# Hydraulic diversity of El Cien Formation (Baja California Sur, Mexico) and the consequences of functional diversity in paleoclimate estimation using fossil wood

**DOI:** 10.1101/768283

**Authors:** Hugo I. Martínez-Cabrera, Emilio Estrada-Ruiz

## Abstract

Community assembly processes, environmental filtering and limiting similarity, determine functional traits values within communities. Because environment influences the number of viable functional strategies species might take, a strong effect of environmental filter often results in communities having species with similar trait values and narrow functional niches. On the other hand, limiting similarity lead to communities with broader functional spaces. The degree to community assembly processes influence wood trait variation has important implications for paleoclimate estimation using fossil wood since the main tenet of the approach is environmental driven trait convergence, and assumes a central role of environmental filtering. We used functional diversity (FD) to determine how three wood anatomical traits vary in 14 extant communities (272 species) growing under different climate regimes, and inferred the prevalence of environmental filtering/limiting similarity. We also calculated FD metrics for the El Cien Formation fossil woods and discussed the results in light of the current knowledge of the flora. We found lower anatomical diversity in communities growing in dry/cool places (smaller functional spaces and lower abundance of trait combinations), suggesting that strong wood anatomical trait convergence could be the result of stronger habitat filtering in these communities. A lower strength of environmental filter in warm/wet environments, likely results in an amplification of the role of other drivers that promote higher number of hydraulic strategies through niche partition in highly structured communities. More complex ecological structures in mild tropical places likely lead to a higher spread of wood trait values. This asymmetry in the strength of environmental filter along climate gradients, suggest that the imbalances in strength of the trait-climate convergence, should be incorporated in paleoclimate prediction models. FD approach can be used to recognize promising traits with narrow niches along climate gradients, and therefore a constant effect of environmental filter.

## INTRODUCTION

Limited distribution of species in relation to environment is explained by community assembly hypothesis throughout a series of processes occurring at local scale that allow some species, but not others, to coexist (Diamond 1975; Cornwell and Ackerly 2009). These community assembly processes (i.e. environmental filtering and limiting similarity) determine functional trait values within communities (Weiher et al. 1998; Grime 2006; Cornwell and Ackerly 2009). While environmental filter sets limits to the range of ecological strategies and trait values that species might take in particular sites (Weiher and Keddy 1995; Diaz et al. 1998; Cornwell and Ackerly 2009), limiting similarity (particularly competition) constraints the degree of trait similarity among species (MacArthur and Levins 1967). That is, while environmental filter tends to drive trait to a particular value, limiting similarity tends to constrain the degree of trait similarity among species within communities. Environmental filtering occurs when there is high selection pressure on traits. It occurs, for example, when species with very low resistance to drought-induced embolism are excluded from places experiencing drought, resulting in communities with highly convergent trait values conferring cavitation resistance. Limiting similarity occurs when there is a great overlap in niche requirements among species, resulting in competition and local exclusion from a given community. Alternatively, limiting similarity might lead to species coexistence if there is resource partition or results in segregation in different microsites (Chesson 2000). When resource partition is strong, it might be expected an even spacing of trait values in a given community (Cornwell and Ackerly 2009). This resource partition is present, for example, among species in vertically structured communities, where short shade-tolerant species of the understory species tend to be mechanically stronger (Puntz and Brokaw 1989) and higher wood density (van Gelder et al. 2006) than larger light demanding species (Puntz and Brokaw 1989; Falster and Westoby 2005). Along this light gradient, wood anatomical varies with the vertical niche position because vessel size and density are related to plant size (e.g. Preston et al. 2006) to cope with conduction demands associated with increasing height (Ryan and Yoder 1997).

Paleoclimate estimation using fossil wood anatomical traits has largely focused on the effect of the environmental filter on trait values to make predictions (e.g. Weimann et al. 1998, 1999; Martínez-Cabrera and Cevallos-Ferriz 2008). This approach essentially focuses on the strength of environmental filter as it measures the degree of climate convergence of trait values to build their climate prediction models. These models use the relative prevalence of a qualitative trait and/or the difference in trait values under different environments. Limiting similarity, however, might have important consequences in the context of paleoclimate estimation since the possibilities of values a trait might take would increase due to a higher niche diversity in more structured communities, hindering the effect of environmental convergence. That is, while environmental filtering selects for trait convergence (Keddy 1992), species interaction and limiting similarity might have a diversifying effect on trait values (Losos 2008) to allow partition of resources and occupation of a variety of niches. The strength of these two community assembly processes might differ by trait and climate regime, likely having an effect on predictive models of paleoclimate estimation. As a consequence, it would be harder to draw paleoclimatic predictions on highly structured communities compared to those where environmental filter is stronger. It is possible that the prediction models using wood anatomy that have been derived so far have confounding effects since they use traits on which the relative strength of environmental filtering and limiting similarity is still unknown.

In this study, we use a functional diversity framework to determine how wood anatomical trait variation patterns differ in extant communities in relation to the climate they occupy. We particularly focused on hydraulic traits. We tested if communities from mild wet tropical environments have a narrow hydraulic niche variation, which would suggest strong effect environmental filter, or if, on the other hand, despite water availability in these places, the water conduction morphospace is large due to more structured communities in the tropics (stronger effect of limiting similarity). As the functional diversity approach might shed light on the relative strength of the community assembly forces, and ultimately on the general environment, we applied the same framework to describe the different aspects of wood functional trait diversity of El Cien Formation flora to contrast the new evidence with the current ideas of the ecological characteristics of the paleoflora (Martínez-Cabrera and Cevallos-Ferriz 2008; Martinez-Cabrera et al. 2012).

## MATERIAL AND METHODS

We used a previously compiled a dataset of vessel traits from 272 species across 14 communities (Martínez-Cabrera and Cevallos-Ferriz 2008; Schenk et al. 2008; Martínez-Cabrera et al. 2009, 2011, 2018). These plant communities range from tropical rainforest (from Mexico, Brazil and Suriname) to communities with drier climate such as desert vegetation (from southern USA, Mexico and Argentina) and temperate forest (USA) (Table 1).

**Table 1.**
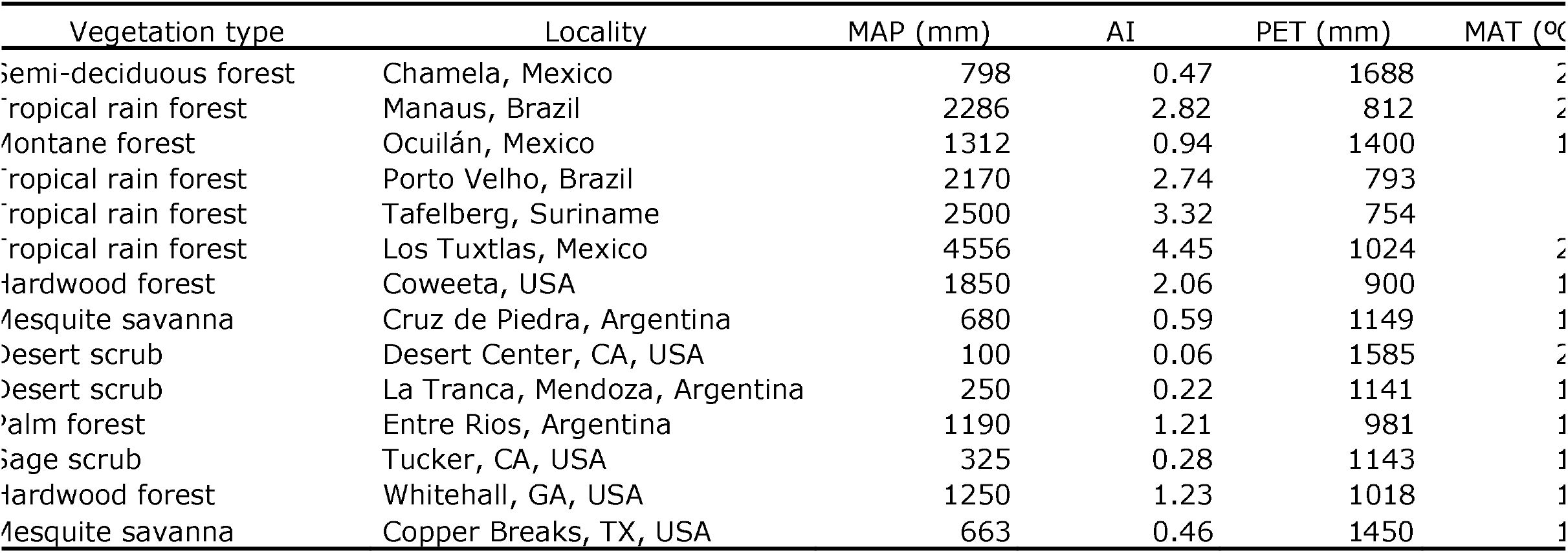
Extant communities and climate variables included in the study.

Vessel lumen diameter was measured on at least 25 random measurements in the tropical communities (Martínez-Cabrera and Cevallos-Ferriz 2008; Martínez-Cabrera et al. 2011). In the communities described in Schenk et al. (2008) and Martínez-Cabrera et al. (2009, 2011), mostly from dry or temperate areas, these traits were measured on 100 to 200 cells. We calculated potential conductivity per stem cross sectional area (*K_s_*) following Zanne et al. (2010): ***K_s_* ∝ *F*^1.5^*S*^0.5^** were *F* is the vessel fraction and *S* is a vessel size contribution metric. Vessel fraction *F* is mean vessel area *Ā* times vessel density (*N*) (***F* − *Ā* N***; mm^2^·mm^-2^), and *S* is the ratio of the same anatomical traits (*S* = *Ā* / *N*; mm^4^). *Ā* is the mean individual vessel cross sectional area and *N* is the vessel number per unit of sapwood area (Zanne et al. 2010). *A* was calculated using mean vessel diameter. *F* is inversely related to mechanical strength (Preston et al. 2006; Zanne et al. 2010), while *S* is directly correlate to a increased cavitation risk that is coupled to high water conduction capacity (Zanne et al. 2010). *F* and *S* represent different axes of variation as they are orthogonal (Zanne et al. 2010).

Climate information, mean annual temperature (MAT), mean annual precipitation (MAP), annual potential evapotranspiration (PET) and aridity index (AI= MAP/PET; aridity index mirrors precipitation, so higher AI is observed in places with high MAP), was obtained from weather stations located in close proximity to the studied localities. For the Mexican montane and tropical rainforest communities we used the ERIC (Extractor Rápido de Información Climatológica) database (INTA 2000). We used the historical records of the weather station at the Biological Station of Chamela (UNAM) to obtain climate information from the Mexican tropical deciduous forest. The climate variables from the Brazilian tropical rainforests communities (Manaus and Porto Velho) were calculated from historical information of a 30-year period (Instituto Nacional do Meteorología, Brazil). For the drier sites in Schenk et al. (2008) and Martínez-Cabrera et al. (2009), climate data was obtained from the nearest weather station with long-term (30-year) climate data.

### Functional diversity metrics

We calculated several complementary functional diversity indices using the vessel traits mentioned above (*K_s_*, *S* and *F*). Functional richness (FRic), functional dispersion (FDis), functional eveness (FEve). FRic measures the amount functional space filled by a community (Villéger et al. 2008), it is the volume in n-dimensional space occupied by species in communities, where n represents the number of traits. In this case, it is the volume occupied by the species in each community in the three dimensions of the measured traits (*K_s_, S* and *F*). We used FRic to determine if the size of the functional space increases, or decreases, with particular climate variables and thus infer if community trait convergence/divergence patterns are related to climate. FDis computes the average distance in trait space of individual species to their group centroid and it is independent of species richness (Laliberté and Legendre 2010). FDis gives information about changes in abundance of trait combinations (Laliberté and Legendre 2009). FEve measures the regularity of spacing between species along functional trait gradients and it is independent of species richness. FEve is 1 when the distances between all nearest neighbor species pairs are identical (Villéger et al. 2008).

We calculated FRic, FDis and FEve using the dbFD function of the FD package (Laliberté et al. 2014) in R (R Core Team 2019). In addition, we used convex hull volumes to visualize the multidimensional trait space of each community. The convex hull is the minimum convex hull that includes all the species in each community. The convex hulls are built by selecting the extreme values for each trait, representing the vertices, to then link them and calculate the volume. The convex hull volume was calculated using the Quickhull algorithm (Barber at al. 1996) as implemented the R package geometry (Roussel et al. 2019).

### El Cien Formation

The El Cien Formation sediments are exposed in the southern part of the Baja California Peninsula (Applegate, 1986), approximately 100 km northwest of La Paz. The fossil woods, preserved as silica permineralizations, were collected in upper member of the formation (Cerro Colorado Member), near Rancho Matanzas, and Cañada El Canelo, about 5 km northeast and 3.5 km southwest from the El Cien, respectively (Fig. 1). The El Cien Formation is a clastic sedimentary sequence deposited during the late Oligocene–early Miocene (Applegate 1986; Fischer et al. 1995). The sediments of the member where the fossil woods were found are mainly composed of fine-to coarse-grained sandstones, tuffaceous sandstones and conglomerates (Applegate 1986; Gidde 1992; Fischer et al. 1995). The current wood diversity of the El Cien Formation is 21 morphotypes, all of them were used for these analyses (Martínez-Cabrera and Cevallos-Ferriz 2008). The samples studied here are housed in National Paleontological Collection, Institute of Geology, Universidad Nacional Autónoma de Mexico (UNAM), Mexico.

**Figure 1.**
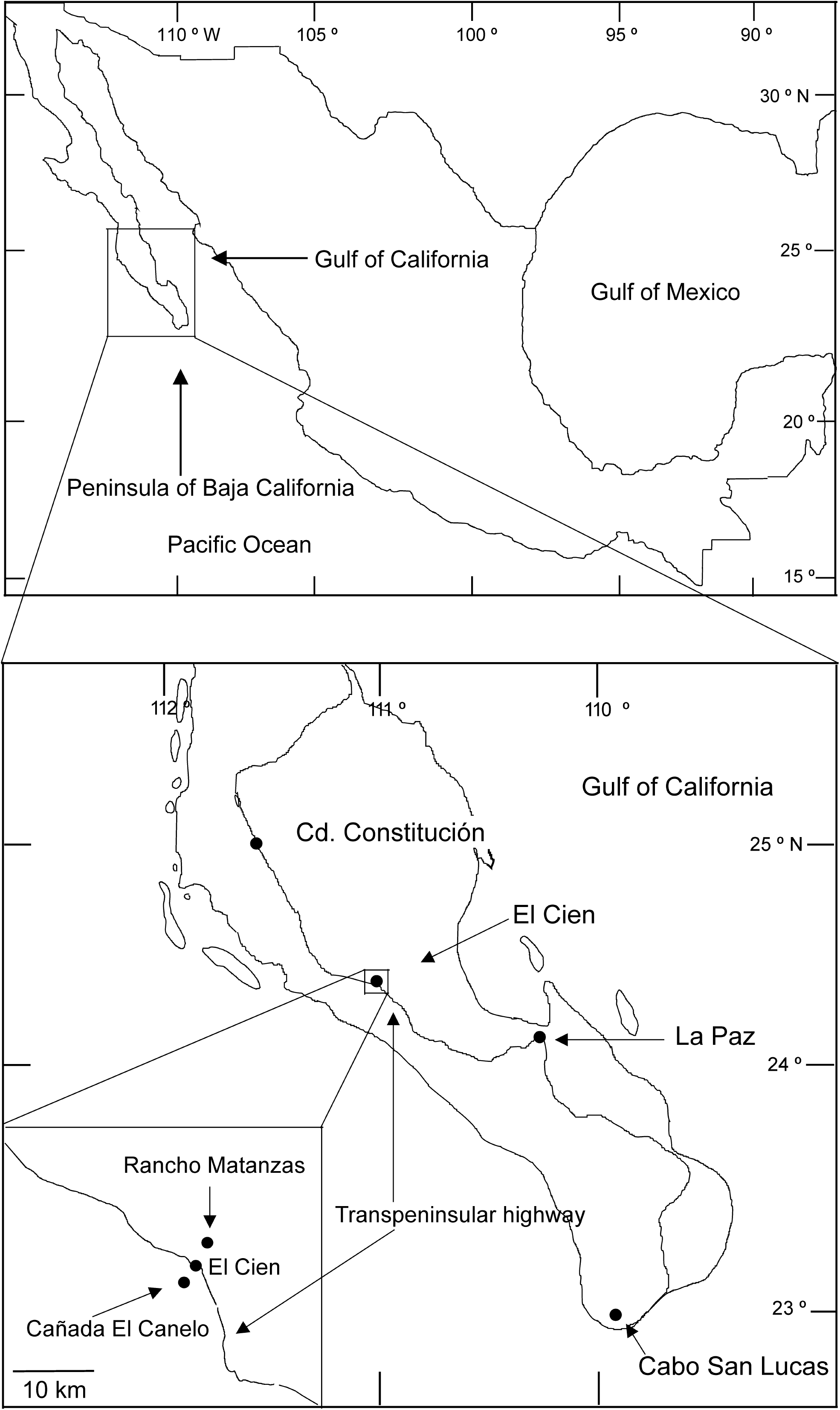
Location map of the El Cien Formation wood flora.

## RESULTS AND DISCUSSION

Functional richness (FRic), the amount of functional space occupied by species in a community is positively related with MAP (r^2^ =0.51, p= 0.003; Fig. 2A), AI (r^2^ =0.40, p= 0.014; Fig. 2B) and MAT (r^2^ =0.39, p= 0.015; Fig. 2C). Therefore, communities in places with higher precipitation, temperature and lower aridity (higher values indicate low aridity) are functional richer (i.e. larger hydraulic niche) than in dry/cold places, where functional convergence results in a more restricted hydraulic niche (Fig. 3A). Functional dispersion (FDis), the overall distance between individual species form the center of the multivariate space, also increases with precipitation (r^2^=0.52, p=0.003; Fig. 2D), AI (r^2^=0.47, p=0.006; Fig. 2E) and temperature (r^2^=0.35, p=0.02; Fig. 2F) indicating that in comparatively colder/ dryer communities trait combinations tend to be more similar among species (there is a lower abundance of trait combinations, Fig. 3B). The regularity of spacing among species (FEve) is not significantly related to any climate variable. PET was not correlated with any of the functional diversity metrics. These results indicate that with higher precipitation, coupled with higher temperature and lower aridity, the hydraulic niche and number of trait combinations is larger than in dry/cool places, suggesting that functional trait convergence is promoted by a stronger effect of the environmental filtering under these climates. However, communities in warm climates might have large hydraulic niches even at relatively low precipitation values (e.g. semi-deciduous forest, Chamela Fig. 4A), while cool temperate communities might have small hydraulic niches even if precipitation is high (hardwood forest, Coweeta, Figs 4A and B). Narrow niches are often associated with a strong habitat filter (e.g., Craven et al. 2018), as it reduces the viable functional strategies of species to face particular environmental pressures (Keddy 1992; Weiher and Keddy 1998). On the other hand, the diversified hydraulic strategies in wet and warm environments might be the result of an increased role of limiting similarity and niche partition, in part because of the relaxation of the abiotic filtering. It is important to mention that limiting similarity and environmental filtering are not competing models of community assembly, as both operate at the same time in a community (Cornwell and Ackely 2009). Cornwell and Ackely (2009) found, evidence of both community assembly processes drive variation on different traits, and in one case (specific leaf area), the effect was simultaneous, with environmental filtering driving mean values along the gradient and intra-community trait value spacing mediated by limiting similarity. Moreover, the relative strength of these two assembly forces could shift in time and with successional position. Craven et al. (2018), for example, found evidence of a greater functional convergence (lower FDis and FRic) in later stages of succession in a seasonally wet tropical secondary forest.

**Figure 2.**
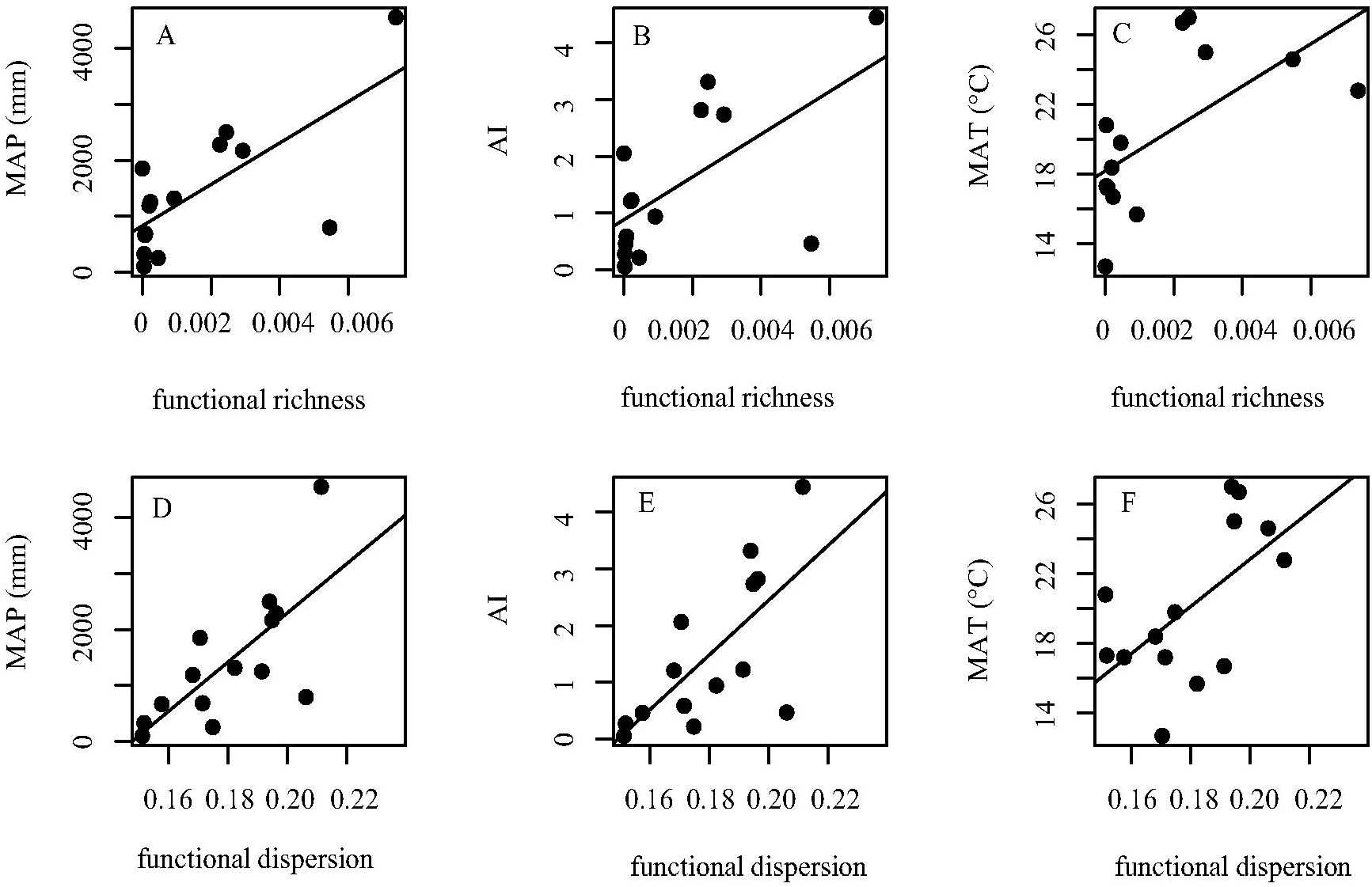
Pair-wise correlations of climate variables with functional richness (Fig. 2 A-C) and functional dispersion (Fig. 2D-2F).

**Figure 3.**
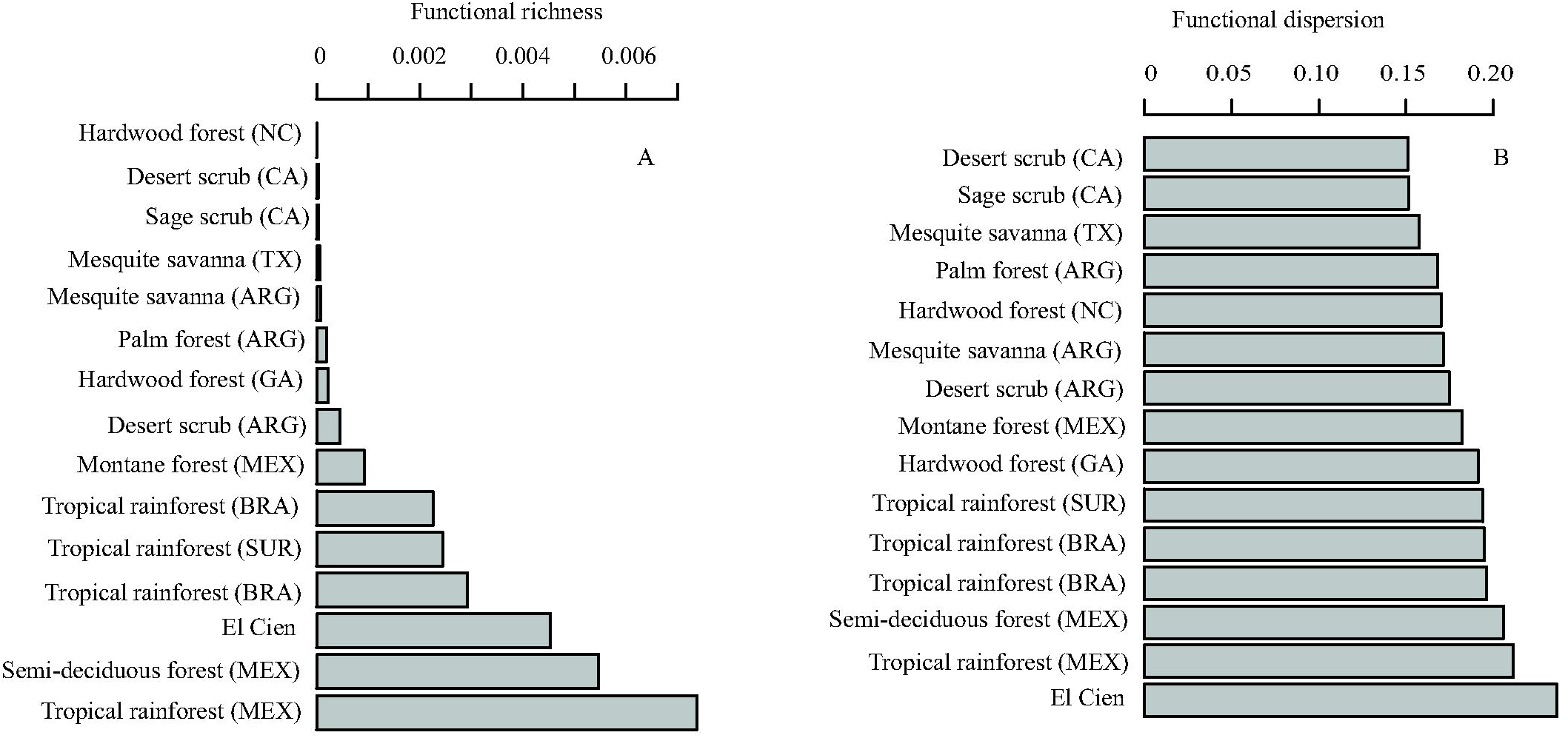
Bargraphs showing functional richness (A) and functional diversity (B) values in extant communities and El Cien Formation wood flora.

**Figure 4.**
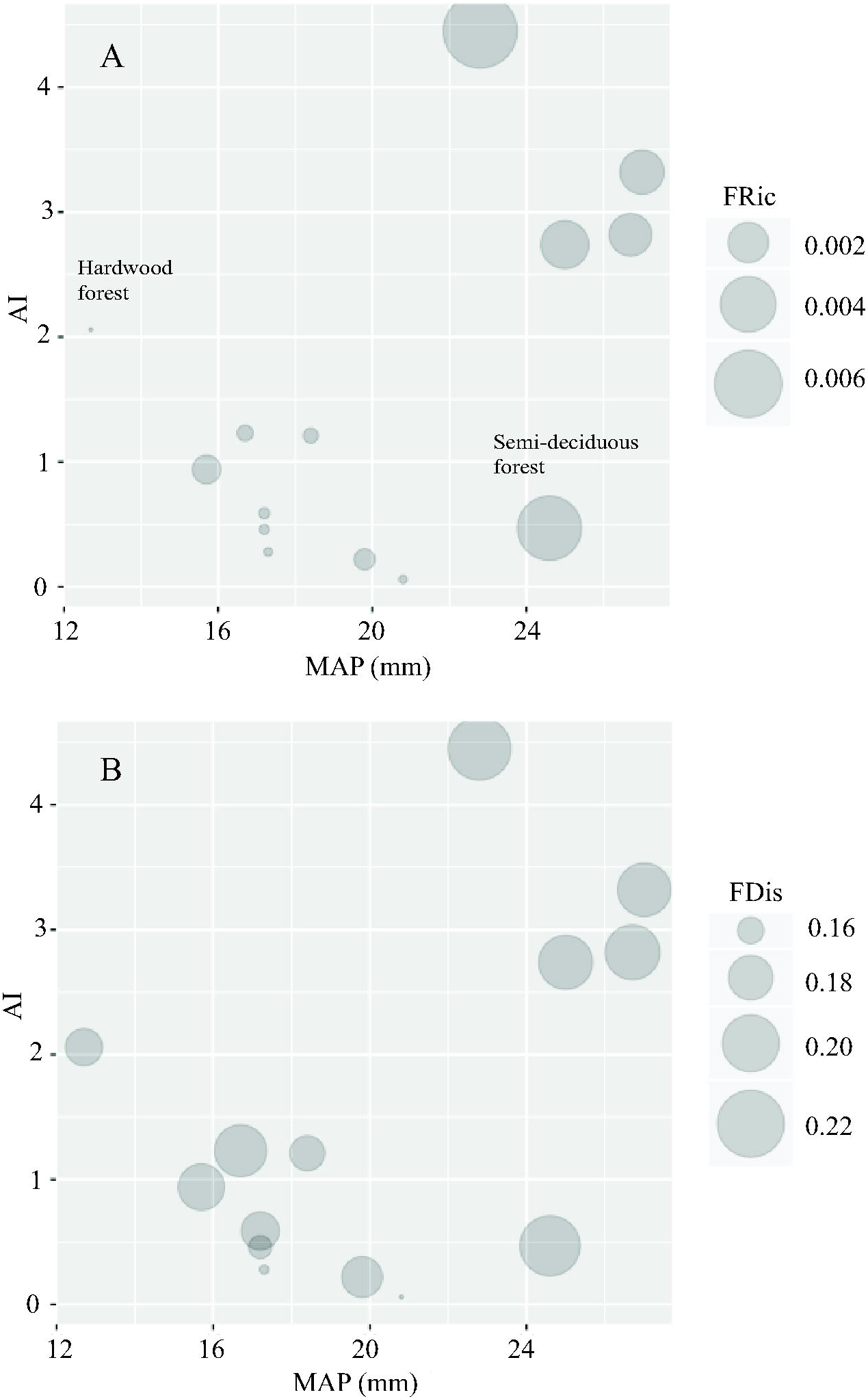
Bubble plots showing the relationship between AI and MAT. The size of the circles represents A) functional richness (FRic) and B) functional dispersion (Fdis). As the relationship of AI and MAP is high, MAP plot is not included.

The results from the convex hull volumes support the functional diversity analyses (Fig. 5). Overall, the volumes of convex hulls of the communities with a wet warmer climate were larger and more complex when compared to those growing in dry/cool habitats (Fig. 5), indicating a higher convergence of hydraulic strategies in the later communities. A more limited multivariate trait space is consistent with strong abiotic filter (Keddy 1992; Cornwell et al. 2006) in these sites (Fig. 5).

**Figure 5.**
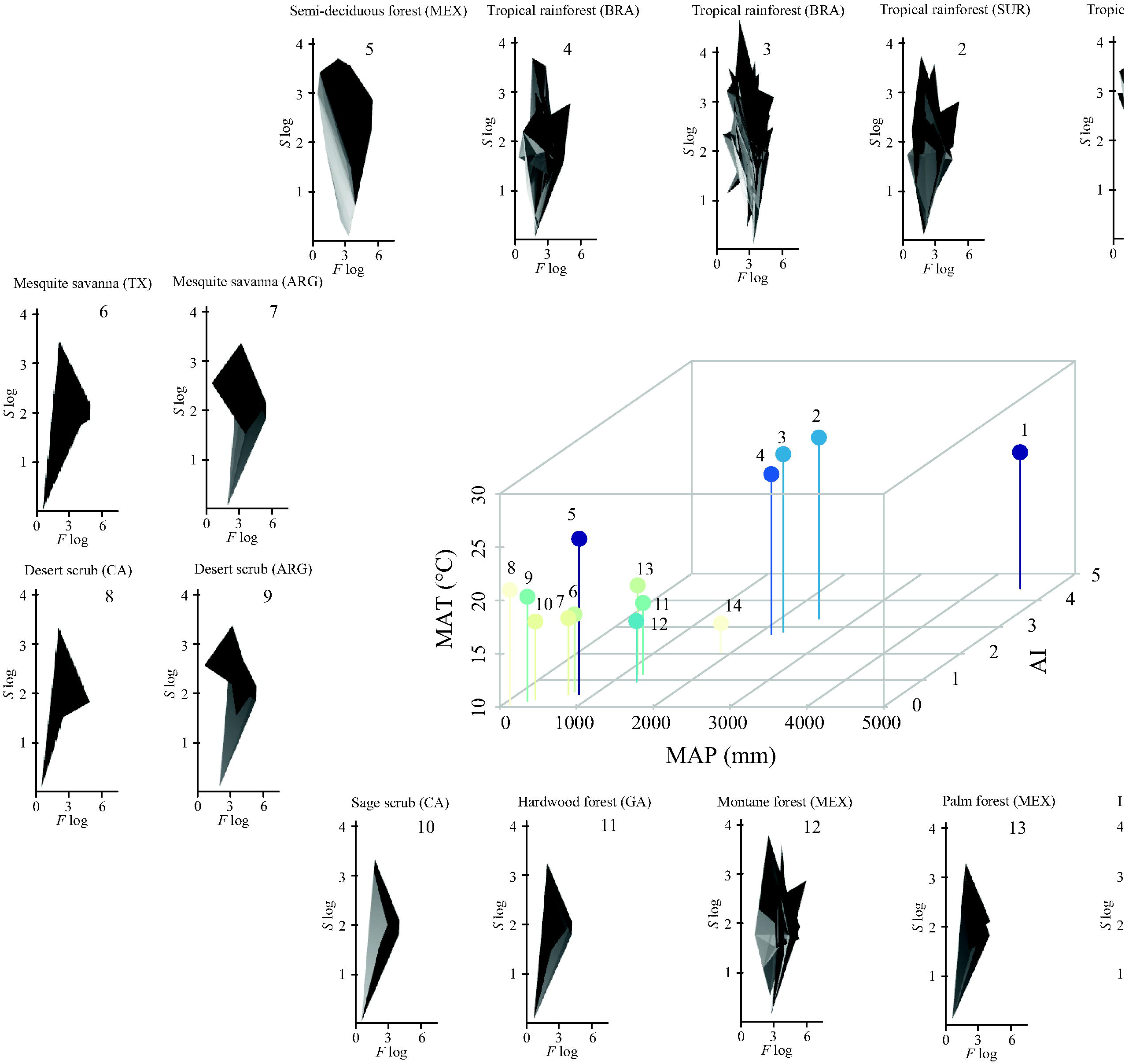
3D Scatter plot of the three climate variables correlated with functional diversity (MAT, MAP and AI) and convex hull volumes for the communities studied. The colors of the circles indicate the magnitude of the FRic, larger in dark blue, smaller in light yellow. Convex hulls indicate the three-dimensional space occupied by all the species in each community; it is a representation of the hydraulic niche. For simplicity, only x (*F* log) and y axes (*S* log) are portrayed, the z-axis (*K_s_* log) was not drawn, but the convex hull are three-dimensional.

### Implications of functional diversity paleoclimate estimation

Predictive paleoclimate models using fossil wood employ trait information of extant communities to build their models. These models assume that environmental trait convergence is constant across communities, and therefore a constant strength of trait-climate convergence. The effect of environmental filtering may, however, limit trait values more strongly in the stressful side of a climate gradient, making these models good predictors on one side of a particular climate gradient, but not on the opposite. In addition, if models were built using communities where the effect of environmental filtering and limiting similarity/niche partition differs, the predictions might have large associated errors. For example, in alpine or boreal forest small vessels are strongly selected as they confer resistance against freezing induced cavitation (e.g., Davis et al. 1999), but on the warmer side of the gradient, temperature selection of vessel size is relaxed, allowing vessel size variation to be driven by water availability (e.g., Hacke et al. 2000), canopy position and succession (e.g., Apgaua et al. 2017), and overall ecological strategy (fast vs. slow growing species; e.g. Apgaua et al. 2017). That is, on one side temperature strongly drives vessel size (small functional niches driven by a strong filter), while on the other the temperature selection on size is relaxed, allowing for larger vessels variation. These results evidenced an asymmetry in the effect of environmental filter on the hydaulic strategies, with smaller hydraulic niches and a lower number of trait combinations in the colder/dryer side of the spectrum, while larger niches are found in the warm/wet side.

Functional diversity analyses can help in paleoclimate estimation by recognizing individual wood traits or multi-trait axes that have narrow variation along particular environmental gradients, where trait convergence operates along the gradient. This approach would also provide additional information on the ecological forces driving trait variation working in paleocommunities.

### Potential of functional diversity in studies of wood anatomical evolution

Evidence suggests that the relationship between wood anatomical traits and climate has not been constant in the history of angiosperms, as some ecologically informative wood traits only appeared in the fossil record until the Paleogene (Wheeler and Bass 1991, 2018). In addition, early in the Cretaceous, there seems to be a low anatomical diversity of angiosperm woods (e.g., Wheeler and Lehman 2009; Martínez-Cabrera et al. 2017) relative to the diversity of other organs (e.g., Crane and Lidgard 1989). During the early Cretaceous (Aptian-Albian) two main fossil wood types dominated the assemblages (*Paraphyllanthoxylon* and *Icacinoxylon/Platanoxylon* e.g. Thayn et al. 1983; Herendeen et al. 1991; Wheeler and Lehman 2009). It is only towards the Cenomanian that there is a relative increase in the frequency of angiosperm fossil woods with a parallel increase of anatomical diversity (Phillipe et al. 2008). Even during the Turonian-Santonian there had been described, only 50 wood types in 2009 (Wheeler and Lehman 2009; Chin et al. 2019). This paucity in angiosperm woods from the early Cretaceous is thought to be the result of preservation (Phillipe et al. 2008), because of the prevalence of thin walls, that was the partial result of the narrow ecological niche they occupied (short plants living in wet, disturbed areas, e.g. Wing and Boucher 1998). Therefore, an integrated approach using phylogenetic and functional diversity methods could inform us if this historical pattern is indeed the results of the environment restricting niche occupancy by imposing a strong filter on traits values and if the subsequent niche diversification, and the increase in morphological disparity (increased divergent evolution among closely related species Martinez-Cabrera et al. 2017) was coupled with an increase in the overall size of the functional landscape or through divergences of closely related species occupying narrow niches.

### Functional diversity of El Cien Formation

The functional space occupied by El Cien Formation woods was among the larger in the communities analyzed here (Fig. 2A). It was only behind the tropical rain forest from Los Tuxtlas and the semi-deciduous forest from Chamela. Surprisingly, the abundance of trait combinations (FDis, Fig. 2B) of El Cien Formation paleocommunity was the largest and similar to floras growing under warmer and humid environments. The hydraulic niche of El Cien flora was large (Fig. 3A, 5). This indicates that the flora had la large hydraulic niche, probably due to the combined effects of a low strength of the environmental filter and a strong effect of limiting similarity that promoted niche partition and a diversified array of strategies. This supports the hypothesis of El Cien Formation flora growing under wet and warm environment. Paleoclimate estimates for El Cien based on wood traits suggest a MAT of 22.5 to 24.8. The evidence regarding water availability is still equivocal. Vessel characteristics indicate that the paleoflora had a hydraulic capacity similar to that observed in some Mexican and South American wet forests (Martínez-Cabrera and Cevallos-Ferriz 2008), suggesting a relatively wet, tropical environment. However, the relatively high prevalence of ring and semi-ring porous woods (Martínez-Cabrera and Cevallos-Ferriz 2008) supports the hypothesis of El Cien paleoflora represents a dryer tropical deciduous or semi-deciduous forest. 40% of the nearest living relatives of from El Cien paleoflora are evergreen, while 20% are deciduous. There is also a close compositional similarity, since about half of the nearest living relatives of the fossils from El Cien are found in the deciduous forest from southwestern coasts of Mexico (Martínez-Cabrera and Cevallos-Ferriz 2008) and the two most diverse families in Chamela (Lott and Atkinson 2002), and the El Cien paleoflora (Martínez-Cabrera and Cevallos-Ferriz 2008) are Leguminosae and Euphorbiaceae (Martínez-Cabrera et al. 2006; Martínez-Cabrera and Cevallos-Ferriz 2008). Further evidence of the similarity between El Cien Formation flora and the deciduous forests of the Mexican western coast is the close estimated value of the paleoflora wood density (Martinez-Cabrera et al. 2012), which is a key trait related to many ecological features including life history strategies, physiological and mechanical traits (e.g., van Gelden et al. 2006; King et al. 2006; Chave et al. 2009).

The functional diversity analyses presented here rather than inform us on the magnitude of a particular climate variable in fossil flora, they provide hints on the particular assembly processes driving their trait variation. The hydraulic diversity of El Cien was large, likely indicating a highly structured niche and therefore prominent role of limiting similarity. In general, higher environmental stress limits trait variation (e.g. wood density, Swenson and Enquist 2007), so that the stronger the environmental limitations the higher the trait convergence. As in milder environments this restriction to trait variation is relaxed, larger functional diversity is observed as a consequence of niche partition in highly structured communities (Muller-Landau 2004).

## CONCLUSIONS

Communities in wet/warm climates in general had higher hydraulic diversity than communities growing in dry/cool environments indicating the presence of a strong environmental filter on water conduction cells in the latter environments. In warmer/ wetter environments strong selection of small sized vessel is relaxed and higher functional diversity is observed, likely due to niche partition present in more structurally complex communities. Our results suggest that more complex ecological structures in mild tropical places possibly lead to a higher spread of wood trait values, making climate prediction using fossil wood more difficult in these biomes. It is necessary to incorporate the effect of asymmetrical environmental convergence in paleoclimate estimation models that use wood anatomical information. Functional diversity could be used to identify traits with constant environmental convergence along climate gradients.

## ACKNOWLEDMENTS

We thank Deborah Woodcock for her comments on a previous version of the manuscript.

## REFERENCES

Apgaua, DM, Tng DY, Cernusak LA, Cheesman AW, Santos RM, Edwards WJ, Laurance SG. 2017. Plant functional groups within a tropical forest exhibit different wood functional anatomy. Funct. Ecol. 31: 582–591.

Applegate SP. 1986. The El Cien Formation, strata of Oligocene and early Miocene age in Baja California Sur. Revista del Instituto de Geología de la Universidad Nacional Autónoma de Mexico 6: 145–162.

Barber CB, Dobkin DP, Huhdanpaa HT. 1996. The Quickhull algorithm for convex hulls. ACM Transactions on Mathematical Software 22: 469–483.

Chin K, Estrada-Ruiz E, Wheeler EA, Upchurch, GR, Wolfe DG. 2019. Early angiosperm woods from the mid-Cretaceous (Turonian) of New Mexico, USA: *Paraphyllanthoxylon,* two new taxa, and unusual preservation. Cretaceous Research 261: 292–304.

Chesson PL. 2000. Mechanism of maintenance of species diversity. Annu. Rev. Ecol. Systemat. 3: 343–366.

Chave J, Andalo C, Brown S, Cairns MA, Chambers JQ, Eamus D, Fölster H, Fromard F, Higuchi N, Kira T, Lescure JP, Nelson BW, Ogawa H, Puig H, Riéra B, Yamakura T. 2005. Tree allometry and improved estimation of carbon stocks and balance in tropical forests. Oecologia 145: 87–99.

Cornwell WK, Schwilk DW, Ackerly DD. 2006. A Trait based test for habitat filtering: convex hull volume. Ecology 87: 1465–1471.

Cornwell WK, DD Ackerly. 2009. Community assembly and shifts in plant trait distributions across an environmental gradient in coastal California. Ecol. Monogr. 79: 109–126.

Crane PR, Lidgard S. 1989. Angiosperm diversification and paleolatitudinal gradients in Cretaceous floristic diversity. Science 246: 675–678.

Craven D, Hall JS, Berlyn GP, Ashton MS, Van Breugel M. 2017. Environmental filtering limits functional diversity during succession in a seasonally wet tropical secondary forest. J. Veg. Sci. 29: 511–520.

Davis SD, Sperry JS, Hacke UG. 1999. The relationship between xylem conduit diameter and cavitation caused by freezing. Am. J. Bot. 86: 1367–1372.

Diamond JM. 1975. Assembly of species communities. Pages 342–444 *in* ML Cody, JM Diamond eds. Ecology and evolution of communities. Harvard University Press, Cambridge, Massachusetts, USA.

Diaz S, Cabido M, Casanoves F. 1998. Plant functional traits and environmental filters at a regional scale. J. Veg. Sci. 9: 113–122.

Falster DS, Westoby M. 2005. Alternative height strategies among 45 dicot rain forest species from tropical Queensland. Aust. J. Ecol. 93: 521–535.

Fischer R, Glli-Olivier C, Gidde A, Schwennicke T. 1995. The El Cien Formation of southern Baja California, Mexico: stratigraphic precisions. Newsl. Stratigr. 32: 137–161.

Freschet GT, Dias AT, Ackerly DD, Aerts R, van Bodegom PM, Cornwell WK, Dong M, Kurokawa H, Liu G, Onipchenko VG, Ordoñez JC, Peltzer DA, Richardson SJ, Shidakov II, Soudzilovskaia NA, Tao J, Cornelissen JH. 2011. Global to community scale differences in the prevalence of convergent over divergent leaf trait distributions in plant assemblages. Glob. Ecol. Biogeogr. 20: 755–765.

Grime JP. 2006. Trait convergence and trait divergence in herbaceous plant communities: mechanisms and consequences. J. Veget. Sci. 17: 255–260.

Gidde A. 1992. Sedimentology of the Miocene Cerro Colorado Member (upper part of the El Cien Formation in Baja California Sur, Mexico). Zbl. Geol. Palaont. 6 (Teil I): 1467–1477.

Hacke UG, Sperry JS, Pittermann J. 2000. Drought experience and cavitation resistance in six shrubs from the Great Basin, Utah. Basic and Applied Ecology 1: 31–41.

Herendeen PS. 1991. Lauraceous wood fromthemid-Cretaceous Potomac group of eastern North America: *Paraphyllanthoxylon marylandense* sp. nov. Rev. Palaeobot. Palynol. 69: 277–290.

INTA 2000. Extractor rápido de información climatológica. Instituto Nacional de Tecnología del Agua, México (ERIC).

Keddy PA. 1992. Assembly and response rules—2 goals for predictive community ecology. J. Veg. Sci. 3:157–164.

King DA, Davies SJ, Tan S, Nur Supardi MN. 2006. The role of stem density and stem support cost in the growth and mortality of tropical trees. J. Ecol. 94: 670–680.

Laliberté E, Legendre P, Shipley B. 2014. FD: measuring functional diversity from multiple traits, and other tools for functional ecology. R package version 1.0-12.

Laliberté E, Legendre P. 2010. A distance-based framework for measuring functional diversity from multiple traits. Ecology 91: 299–305.

Losos JB. 2008. Phylogenetic niche conservatism, phylogenetic signal and the Relationship between phylogenetic relatedness and ecological similarity among species. Ecology Letters 11: 995–1007.

Lott EJ, Atkinson TH. 2002. Biodiversidad y fìtogeografía de Chamela-Cuixmala, Jalisco. Pages 83–97 *In* FN Noguera, JH Vega Rivera, AN García Alderete, M Quesada Avendaño, eds. Historia Natural de Chamela. Inst. Biol., Universidad Nacional Autónoma de México, Mexico.

MacArthur R, Levins R. 1967. The limiting similarity, convergence, and divergence of coexisting species. American Naturalist 101: 377–385.

Martínez-Cabrera HI, Cevallos-Ferriz SRS, Poole I. 2006. Fossil woods from early Miocene sediments of the El Cien Formation, Baja California Sur, Mexico. Rev. Palaeobot. Palyno. 138: 141–163.

Martínez-Cabrera HI, Cevallos-Ferriz SRS. 2008. Palaeoecology of the Miocene El Cien Formation (Mexico) as determined from wood anatomical characters. Rev. Palaeobot. Palyno. 150: 154–167.

Martínez-Cabrera HI, Jones CS, Espino S, Schenk HJ. 2009. Wood anatomy and wood density in shrubs: responses to varying aridity along transcontinental transects. Am. J. Bot. 96: 1388–1398.

Martínez-Cabrera HI, Schenk HJ, Cevallos-Ferriz SRS, Jones CJ. 2011. Integration of vessel trait, wood traits and height in angiosperm shrubs and trees. Am. J. Bot. 98: 915–922.

Martínez-Cabrera HI, Estrada-Ruiz E, Castañeda-Posadas C, Woodcock D. 2012. Wood specific gravity estimation based on wood anatomical traits: Inference of key ecological characteristics in fossil assemblages. Rev. Palaeobot. Palyno. 187: 1–10.

Martínez-Cabrera HI, Zheng J, Estrada-Ruiz E. 2017. Wood functional disparity lags behind taxonomic diversification in angiosperms. Rev. Palaeobot. Palyno. 246: 251–257.

Martínez-Cabrera HI, Estrada-Ruiz E. 2018 Influence of phylogenetic relatedness on paleoclimate estimation using fossil wood: Vessel and fiber-related traits. Rev. Palaeobot. Palyno. 251: 73–77.

Muller-Landau HC. 2004. Interspecific and intersite variation in wood specific gravity of tropical trees. Biotropica 36: 20–32.

Philippe M, Gomez B, Girard V, Coiffard C, Daviero-Gomez V, Thevenard F, Billon-Bruyat J-P, Guiomar M, Latil J-L, Le loeuff J, Néraudeau D, Olivero D, Schlögl J. 2008. Woody or not woody? Evidence for early angiosperm habit from the Early Cretaceous fossil wood record of Europe. Palaeoworld 17: 142–152.

Preston KA, Cornwell WK, DeNoyer JL. 2006. Wood density and vessel traits as distinct correlates of ecological strategy in 51 California coast range angiosperms. New Phytologist 170: 807–818.

Putz FE, Brokaw NVL. 1989 Sprouting of broken trees on Barro Colorado Island, Panama. Ecology 70: 508–512.

R Core Team. 2019. R: A language and environmetn for statistical computing. R Foundation for Statistical Computing, Viena, Austria. URL http://www.R-project.org/.

Roussel JR, Barber CB, Habel K, Grasman R, Gramacy RB, Mozharovskyi P, Sterratt DC. 2019. Package Geometry.

Ryan GM, Yoder BJ. 1997. Hydraulic limits to tree height and tree growth. BioScience 47: 235–242.

Schenk HJ, Espino S, Goedhart CM, Nordenstahl M, Martínez-Cabrera HI, Jones CS. 2008. Hydraulic integration and shrub growth form linked across continental aridity gradients. Proc. Natl. Acad. Sci. 105: 11248–11253.

Swenson NG, Enquist BJ. 2007. Ecological and evolutionary determinants of a key plant functional trait: wood density and its community wide variation across latitude and elevation. Am. J. Bot. 94: 451–459.

Thayn GF, Tidwell WD, Stokes WL. 1983. Flora of the Lower Cretaceous Cedar Mountain Formation of Utah and Colorado. Part 1: *Paraphyllothoxylon utahense*. Great Basin Nat 43: 394–402.

van Gelder HA, Poorter L, Sterck FJ. 2006. Wood mechanics, allometry, and life-history variation in a tropical rain forest tree community. New Phytologist 171: 367–378.

Villéger S, Mason NWH, Mouillot D. 2008. New multidimensional functional diversity indices for a multifaceted framework in functional ecology. Ecology 89: 2290–2301.

Weiher E, Keddy PA. 1999. Ecological assembly rules: perspectives, advances, retreats. Cambridge University Press, Cambridge, UK.

Weiher E, Clarke GDP, Keddy PA. 1998. Community assembly rules, morphological dispersion, and the coexistence of plant species. Oikos 81: 309–322.

Wiemann MC, Wheeler EA, Manchester S, Portier KM. 1998. Dicotyledonous wood anatomical characters as predictor of climates. Palaeogeogr. Palaeoclimatol. Palaeoecol. 139: 83–100.

Wiemann MC, Manchester S, Wheeler EA. 1999. Paleotemperature estimation from dicotyledonous wood anatomical characters. Palaios 14: 459–474.

Wing SL, Boucher LD. 1998. Ecological aspects of the Cretaceous flowering plant radiation. Annu Rev Earth Planet Sci 26: 379–421.

Wheeler EA, Baas P. 1991. A survey of the fossil record for dicotyledonous wood and its significance for evolutionary and ecological wood anatomy. IAWA Bull 12: 275–332.

Wheeler EA, Lehman TM. 2009. New Late Cretaceous and Paleocene dicot woods of Big Bend National Park, Texas and review of Cretaceous wood characteristics. IAWA J 30: 293–318.

Wheeler EA, Baas P. 2018. Wood evolution: Baileyan trends and Functional traits in the fossil record. IAWA J 2–42.

Zanne, AE, Westoby M, Falster DS, Ackerly DD, Loarie SR, Arnold EJS, Coomes D. 2010. Angiosperm wood structure: global patterns in vessel anatomy and their relation to wood density and potential conductivity. Am. J. Bot. 97: 207–215.

